# Detecting phylodiversity-dependent diversification with a general phylogenetic inference framework

**DOI:** 10.1101/2021.07.01.450729

**Authors:** Francisco Richter, Thijs Janzen, Hanno Hildenbrandt, Ernst C. Wit, Rampal S. Etienne

## Abstract

Diversity-dependent diversification models have been extensively used to study the effect of ecological limits and feedback of community structure on species diversification processes, such as speciation and extinction. Current diversity-dependent diversification models characterise ecological limits by carrying capacities for species richness. Such ecological limits have been justified by niche filling arguments: as species diversity increases, the number of available niches for diversification decreases.

However, as species diversify they may diverge from one another phenotypically, which may open new niches for new species. Alternatively, this phenotypic divergence may not affect the species diversification process or even inhibit further diversification. Hence, it seems natural to explore the consequences of phylogenetic diversity-dependent (or phylodiversity-dependent) diversification. Current likelihood methods for estimating diversity-dependent diversification parameters cannot be used for this, as phylodiversity is continuously changing as time progresses and species form and become extinct.

Here, we present a new method based on Monte Carlo Expectation-Maximization (MCEM), designed to perform statistical inference on a general class of species diversification models and implemented in the R package `emphasis`. We use the method to fit phylodiversity-dependent diversification models to 14 phylogenies, and compare the results to the fit of a richness-dependent diversification model. We find that in a number of phylogenies, phylogenetic divergence indeed spurs speciation even though species richness reduces it. Not only do we thus shine a new light on diversity-dependent diversification, we also argue that our inference framework can handle a large class of diversification models for which currently no inference method exists.

## 1 Introduction

The hypothesis of diversity-dependent diversification posits that diversification processes at macro-evolutionary scales are affected by community structure, and particularly by diversity [Walker and Valentine, 1984, Gould et al., 1977]. One of the underlying ideas is that there are ecological limits to diversity (there is a limited number of niches that can be filled with species) and hence to diversification [Rabosky, 2009]. The hypothesis has been extensively studied both empirically and theoretically [Condamine, 2018, Gibb et al., 2016, Cunha et al., 2017, Pouchon et al., 2018, Chen et al., 2017, Pinto-Ledezma et al., 2017, McGuire et al., 2014, Pyron and Wiens, 2013, Xu and Etienne, 2018, Etienne et al., 2016, Liow et al., 2010, Herrera-Alsina et al., 2018, Morlon, 2014, Rabosky and Hurlbert, 2015, Jønsson et al., 2012]. However, currently developed inference models for detecting diversity-dependent diversification from molecular phylogenies consider only species richness as a proxy for diversity [Etienne et al., 2012a].

Phylogenetic diversity, quantifying the genetic differences among a group of species, has been identified as a key feature of diversity [Kling et al., 2018, Scheiner et al., 2017] to be taken into account in conservation biology [Laity et al., 2015, Faith and Baker, 2006] (but see Cantalapiedra et al. [2019], Mazel et al. [2018]), community ecology [Stadler et al., 2017, Tucker et al., 2016, Webb et al., 2006, Violle et al., 2011], evolutionary biology [Kling et al., 2018] and the intersection of these fields. Phylogenetic diversity, or phylodiversity, provides a different perspective on diversity and ecological limits. On the one hand, species richness models suggest that as species diverge there may be less opportunity to speciate further because the growing phenotypic space between species leaves less room to be occupied. However, one could argue that on the other hand, the divergence provides access to increased phenotypic space to speciate into. Hence, extending diversity-dependence to phylodiversity and developing methods to infer such phylodiversity-dependent diversification from molecular phylogenies seems worthwhile. It will allow us to consider the dynamic nature of ecological limits [Costa et al., 2008, Lister, 1976, Soininen et al., 2011] and thus relax the assumption of fixed limits [Marshall and Quental, 2016, Etienne et al., 2012a].

The incorporation of phylogenetic diversity is not possible with the current simulation-free methods for inferring diversity-dependent diversification using the Q-approach introduced by Etienne et al. [2012a] and Laudanno et al. [2020] and implemented in the R package `DDD`. The Q-approach is based on a hidden Markov model approach where the probability of an extant-species tree is integrated over the infinite set of complete trees compatible with it, i.e., the trees that also contain now-extinct species. By only taking into account the number of species at any point in time, the Q-approach does not depend on tree topology. Contrastingly, phylogenetic diversity, defined as the sum of the lengths of all branches in a phylogenetic tree [Faith, 1992], highly depends on the topology of the tree as well as the branching times. Hence, a new methodology is needed to incorporate topological characteristics of the diversification processes.

To do so we generalize a recently developed statistical framework [Richter et al., 2020] based on Monte Carlo Expectation-Maximization (MCEM) that allows inference on a general class of diversification models, including models with phylodiversity-dependent diversification. In this EMPHASIS (Expectation-Maximization in PHylogenetic Analysis with Simulations and Importance Sampling) framework, maximum likelihood estimation is performed on an augmented data set generated by Monte Carlo simulations.

The general class of Species Diversification Models (SDM) introduced in Richter et al. [2020] contains a broad spectrum of scenarios considered in the literature where rates can be constant [Nee et al., 1994], related to the age of the species [Hagen et al., 2015], to the (changing) paleo-enviroment [Descombes et al., 2018], to geographic patterns [Goldberg et al., 2011] or to temperature and diversity [Condamine et al., 2019], just to name a few. For each of these models specific likelihood formulas have been derived (implemented in different packages), but our new method can handle all of these within a single framework, and also applies to combinations of these models for which no such likelihood formula is available and is often impossible to derive or compute numerically. It also applies to new models such as the phylodiversity-dependent models discussed in detail here, and other models with possibly complex interactions between ecological factors and macroevolution, thereby opening endless opportunities for macroevolutionary diversification analysis. The main challenge of our framework is computationally: the Monte Carlo integration is very demanding. In this manuscript we therefore provide a method to perform this integration efficiently.

We illustrate our inference method by applying it to 14 phylogenies, comparing a phylodiversity-dependent diversification model to a diversity-dependent diversification model. We generally find little difference between these two models, although the phylodiversity-dependent diversification model provides an additional narrative for the evolution of global speciation through time in several cases.

## 2 Diversity-Dependent Diversification Models

Diversity-dependent species diversification models are typically used to quantify the effect that diversity has on diversification [Etienne et al., 2012b, Cunha et al., 2017, Etienne and Haegeman, 2012, Foote et al., 2018, Condamine et al., 2019]. Under the classical linear diversity-dependent diversification (LDD) model, it is assumed that speciation rate is a linear function of species richness:

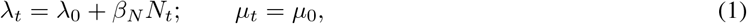

where *λ*_*t*_ is the per species speciation rate, *N*_*t*_ the number of species and *µ*_*t*_ represents the per species extinction rate, at time *t*. Assuming that *λ*_0_ is positive, if *β*_*N*_ is negative the quantity *K′* = − *λ*_0_*β*_*N*_ is called the carrying capacity, which denotes the value for which a clade approaches a niche limit and consequently experiences a slow-down in speciation. If *β*_*N*_ = 0, then the model reduces to the diversity-independent diversification model, i.e. the constant-rate model.

Phylogenetic diversity is recognised as a critical feature to take into consideration in several fields such as conservation ecology, macroecology and macroevolution. So far, it has been studied mostly in a qualitative way and only as a single number at the present instead of considering it as a dynamical quantity that changes through macroevolutionary time. Current diversity-dependent diversification models do not consider phylodiversity, and assume that diversity slows down diversification (e.g. due to niche filling), while qualitative studies suggest that diversity can spur diversification [Jarne et al., 2017, Hamilton et al., 2020]. Likelihood-based inference approaches ignore phylodiversity; they fully describe the processes by considering the probability that the clade has *N*_*t*_ lineages at time *t*, but ignore the topology of the trees. We here introduce a generalised diversity-dependent diversification model, i.e., a phylodiversity-dependent diversification (LPD) model, where we assume that the speciation rate also depends on the phylogenetic diversity per species:

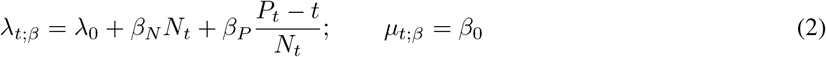

where *P*_*t*_ is the phylogenetic diversity at time *t* defined as the total branch length of the reconstructed tree up until *t*. The quantity 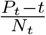 corresponds to the phylogenetic diversity per species at time *t*. Note that by subtracting *t* from the phylogenetic diversity *P*_*t*_, the phylogenetic diversity per species remains 0 for a single species. In Figure 1, an example tree is plotted, with the species richness though time and the phylogenetic diversity per species through time.

**Figure 1:**
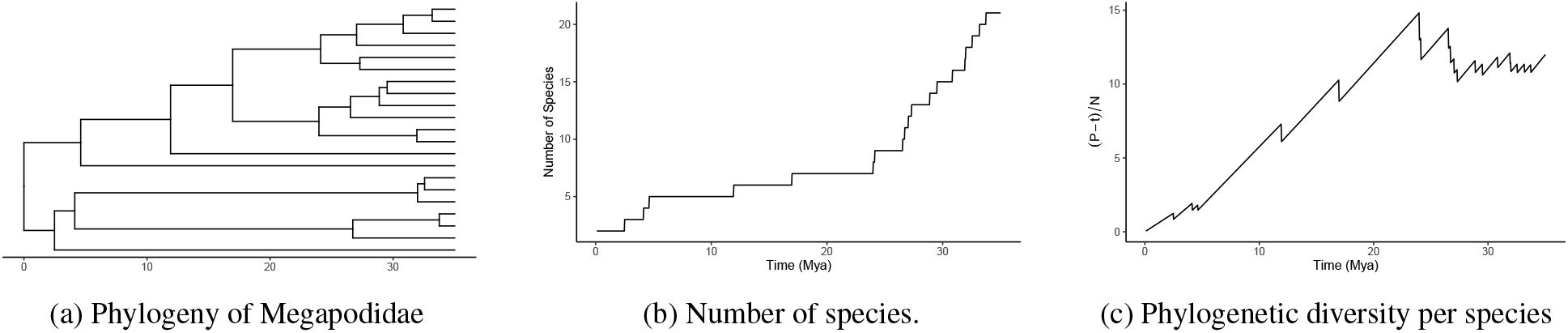
Phylogeny, number of species and phylogenetic diversity per species.

Here, we make use of the statistical methods described in Richter et al. [2020], combined with an efficient importance sampler, in order to perform statistical inference assuming diversification dynamics given by the LPD model and compare it with the diversification dynamics given by the simple LDD model. In this way, we quantify the signal that phylodiversity leaves in species diversification and evaluate if its incorporation in diversity-dependent diversification models is promising for further studies.

## 3 Materials and Methods

Phylogenetic trees are branching diagrams, reconstructed from DNA sequences, representing the evolutionary history of species diversification [Kapli et al., 2020]. Mathematically, they are represented by a discrete part given by the topology of the tree and a continuous part given by its branching times. We define a tree *x* = {**t**, *τ*} as a combination of branching times and topology. More precisely, the branching times are defined by a chronological sequence vector of times **t** = {*t*_0_, *t*_1_, *t*_2_, …, *t*_*p*_}, with *t*_0_ = 0 and *t*_*p*_ being the present time. The topology is defined by a succession of allocation values which can be characterized in multiple ways such as in network or matrix notation. Here, we consider the succession of species names 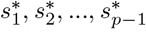 to be the species that diversified (or became extinct) at branching time *t*_*i*_. Moreover, we define the subsets of subindex 𝒞_*x*_ ⊂ {1, …, *p* − 1} and *ε*_*x*_ ⊂ {1, …, *p* − 1} to be the indices corresponding to speciation and extinction events, respectively. This means that if *i* ∈ 𝒞_*x*_ then *t*_*i*_ is a branching time corresponding to a speciation event while if *i* ∈ *ε*_*x*_ then *t*_*i*_ is a branching time corresponding to an extinction event.

Figure 2 shows a representation of a tree describing a full evolutionary process (speciation and extinction events), and the corresponding reconstructed tree, considering the evolutionary history of currently extant species. In ths case 𝒞_*x*_ = {0, 1, 2, 4} and *ε*_*x*_ = {3, 5}.

**Figure 2:**
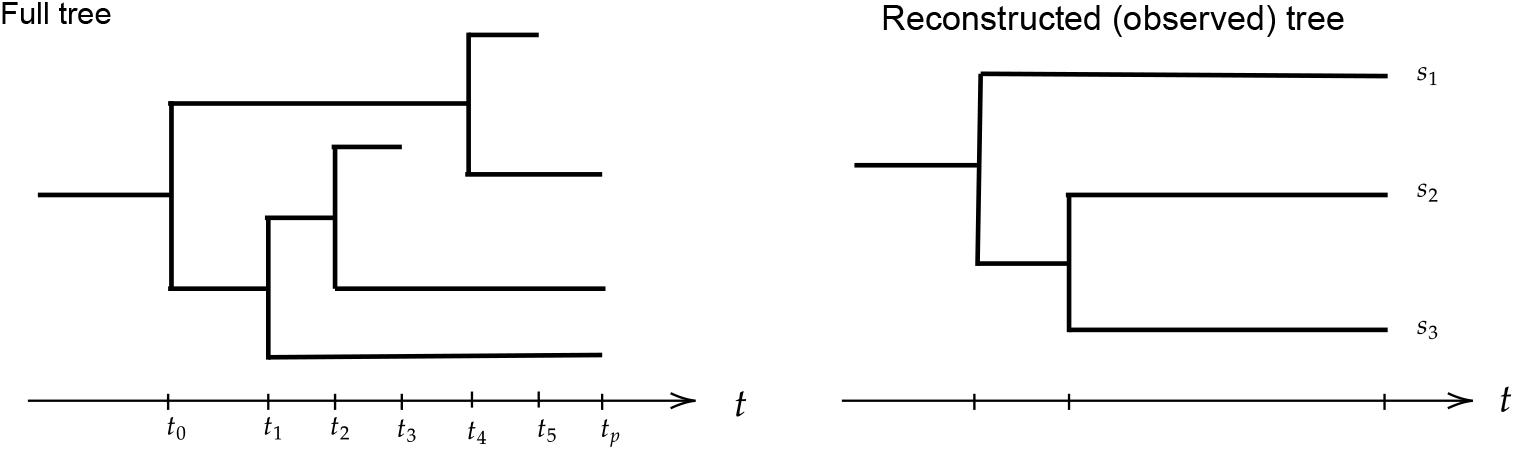
Full phylogenetic trees (left) and the corresponding reconstructed tree (right). Each branch represents a species.

**Figure 3:**
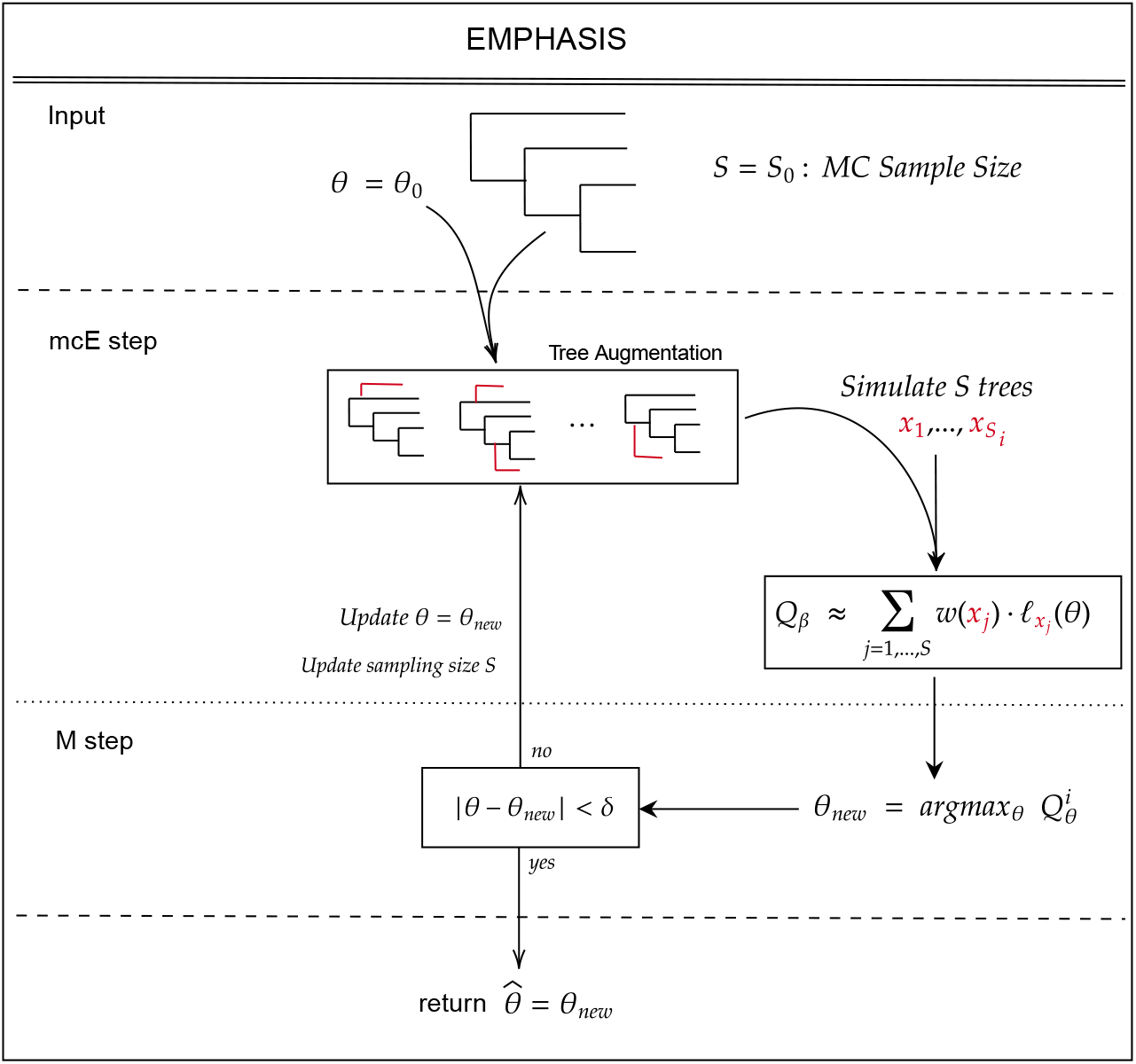
Monte-Carlo EM algorithm diagram in the context of phylogenetic trees.

Throughout, we consider the extant species trees to be accurate (i.e., no uncertainty in branching times or topology). Statistically, extant species trees are our observed data and extinct species are usually latent or unobserved variables, which in the case of diversity-dependent diversification also affect diversification rates.

### 3.1 Diversification of species as a point process

We consider the species diversification process as a general Point Process where each species has a waiting time to speciate into two daughter species that follows an exponential probability distribution with rate *λ*_*t,s*|*θ*_, for any time *t*, species *s* and parameters *θ*. Species can also become extinct with an exponential distribution with rate *µ*_*t,s*|*θ*_ for the waiting time to extinction. We will denote the set of extant species at time *t* by 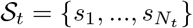, and the number of extant species at time *t* by *N*_*t*_. These quantities are described by a Non-Homogenous Poisson Process (NHPP) [Daley and Vere-Jones, 2007]. Typically, we consider

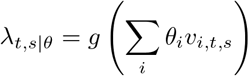

for a set of covariates *υ*_*i,t,s*_ and a link function *g* : ℝ → ℝ [Dobson and Barnett, 2008]. The loglikelihood function of the full process represented by a complete tree (Figure 2, left) is given by

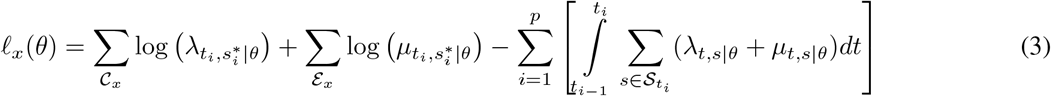

Phylogenies that are derived from molecular data (e.g. DNA sequences) are, however, not full trees, as they do not contain the extinct species (Figure 2 right). The likelihood for an observed tree can be written in terms of the likelihood of compatible full trees. In principle, this is simply the integration over all possible full trees that are in agreement with the observed tree *x*_*obs*_:

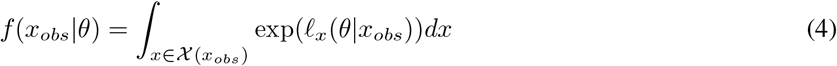

This integration is usually impossible to compute in practice for most diversification models. Here, we present a method where maximum likelihood estimation is possible without calculating directly the likelihood function (4), by implementing a combination of statistical inference and a data augmentation algorithm.

### 3.2 The EMPHASIS Statistical Framework

Our statistical framework is a generalisation of that of Richter et al. [2020], which makes use of an Expectation-Maximization algorithm for maximising the likelihood [Dempster et al., 1977]. The EM algorithm is an iterative procedure consisting of two steps: the E-step and the M-step. Starting from an initial value for the parameters, the E-step involves computing the expected loglikelihood of the observed tree for the given parameters and the M-step involves computing the parameters that maximise that expectation of the loglikelihood. Each iteration the parameters are updated with the values obtained in the M-step of the previous iteration. The E-and M-steps are run iteratively until convergence is reached. The parameters thus obtained have been shown to be the maximum likelihood estimators [Dempster et al., 1977].

Because the expectation in the E-step cannot be computed exactly (or numerically) due to the high dimensionality of the space of complete trees, Richter et al. [2020] proposed to use a stochastic approximation and data augmentation [Tanner and Wong, 1987], specifically a Monte-Carlo method [Chan and Ledolter, 1995] in combination with importance sampling [Glynn and Iglehart, 1989] in the E-step of their EM-algorithm [McLachlan and Krishnan, 2007], and calculated

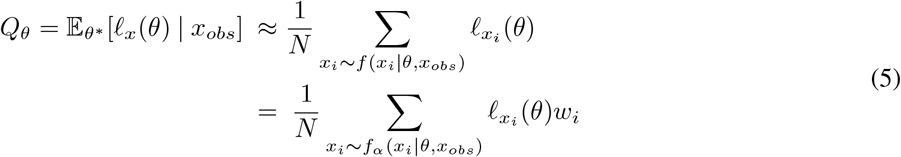

where

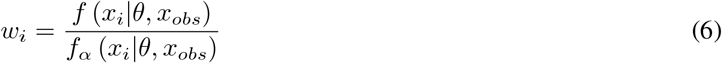

are called the *importance weights* and are highly dependent of the importance sampler *f*_*α*_. The importance weights reflects how accurate the data augmentation is in comparison with the desired distribution, importance weights equal to 1 shows that the importance sampler is the same distribution as the likelihood *f* distribution that generates the process of interest. The data augmentation scheme used by Richter et al. [2020] was mathematically correct but computationally inefficient, as the paper was aimed at the conceptual framework rather than performance. Here we present an improved version of the framework, hereafter called *emphasis*, with a very efficient data augmentation scheme, because the choice of data augmentation scheme is crucial for computational performance [Van Dyk and Meng, 2001].

### 3.3 Augmentation of observed trees, a novel importance sampler for phylogenetic inference

Richter et al. [2020] presents an MCEM algorithm where trees are augmented by drawing uniformly the number of branching events and its corresponding branching times. The method worked well for small trees, but the variance of the estimates grows fast as the tree gets larger for constant number of samples, making the method computationally intractable for medium-sized to large clades. This is due to the curse of dimensionality, i.e., the problem of exploring high-dimensional spaces efficiently [Friedman, 1997]. Our proposed alternative for the data augmentation algorithm augments trees according to the underlying diversification model, encouraging samples in the regions of parameter space that are likely under the proposed SDM.

To sample the extinct species in the tree, we approximate the diversification process of extinct lineages conditional on the extant species in the data as a birth-death process with rates

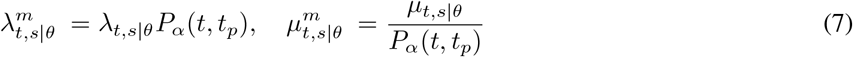

where *P*_*α*_(*t, t*_*p*_) is an approximation of the probability that a species observed at time *t* will not have any descendants at time *t*_*p*_. Kendall [1948] showed that, for lineage-independent models, the exact probability is given by

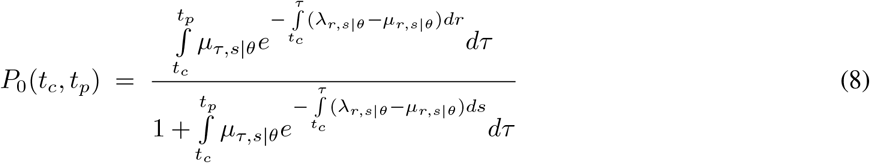

Note that the probability depends on information on *λ*_*ts*|*θ*_ and *µ*_*ts*|*θ*_ for *t*_*c*_ *< t < t*_*p*_. For constant rates this information is available and calculation of Eq. 8 is easy. However, for most SDM information on the full process is not available. For instance, in the case of diversity-dependent diversification models the quantity *N*_*t*_ is unknown.

We augment the observed tree with hidden speciation events. These events can be allocated to all lineages, but not with equal probability. Speciation events occurring on an observed lineage have twice the weight of speciation events occurring on an unobserved lineage [Etienne et al., 2012a]. Figure 4 shows an example when a tree is augmented with a new speciation event at a time that there are two extant lineages and one extinct lineage. In that case, there are five possible allocations. More generally, there are 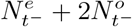 possible allocations, where 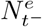 is the number of currently extinct lineages alive just before time *t* and 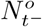 is the number of currently extant lineages just before time *t*. Therefore, we can compute the probability distribution for the waiting times for the augmented speciation events (which we will call missing speciation events) considering the 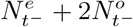 non-homogenous Poisson processes together. Because the minimum waiting time for exponential distributed processes is also an exponential process, given a time *t*_0_, the waiting time for the first missing speciation event to occur is given by an exponential distribution with rate

**Figure 4:**
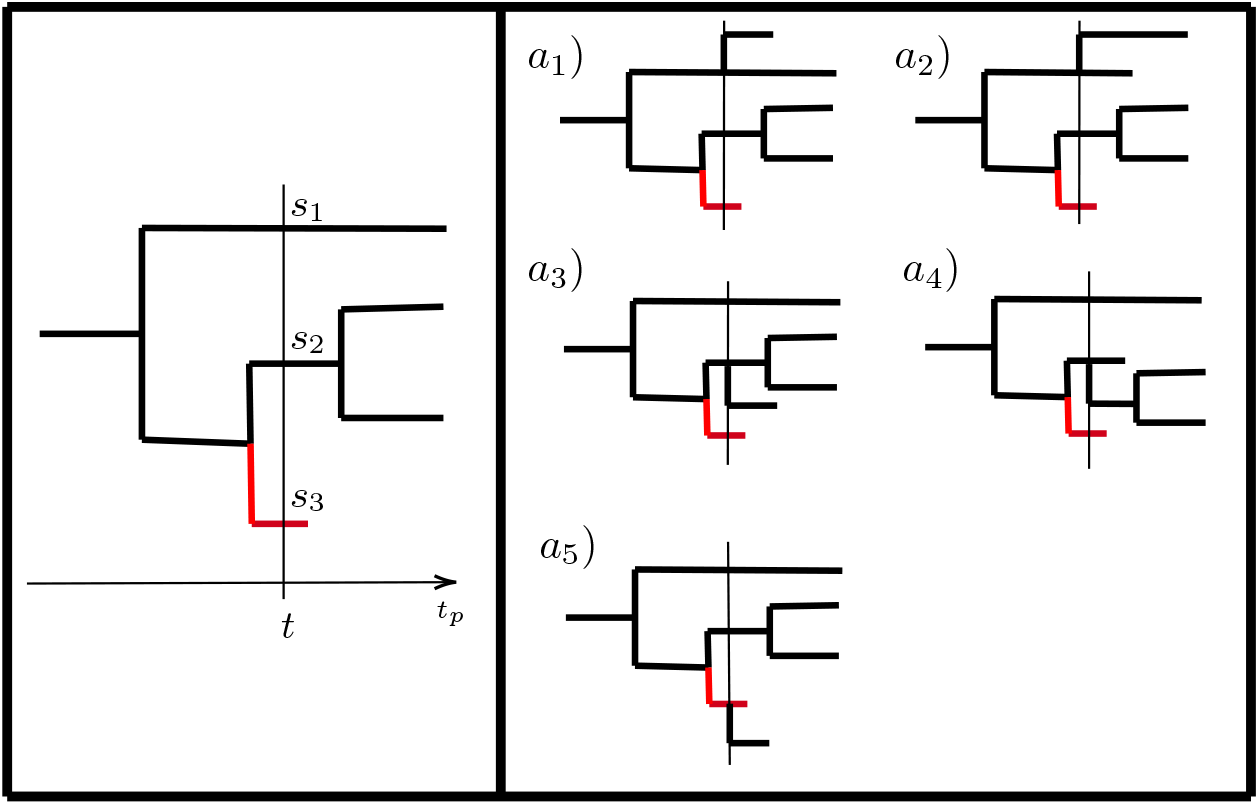
Phylogenetic tree with 2 observed species and 1 missing species at time *t*. When a new species is created there are 2*N*_0_ + *N*_*m*_ possible allocations, in this case 2 ∗ 2 + 1.

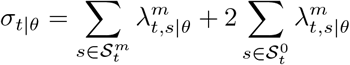

where 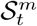 and 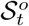 are the sets of observed and missing species at time *t* respectively. Hence, the probability density of the waiting time for any speciation to occur at time *t*, starting the process at initial time *t*_*i*_, is a non-homogeneous exponential distribution with rate *σ*_*t*|*θ*_, that is

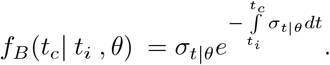

Once a missing speciation event has occurred, the new lineage needs to get an allocation and a extinction time assigned to be included in the tree. In a model where speciation rates are the same for all lineages, all allocations have the same probability

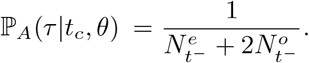

The lineage produced at the missing speciation event must become extinct before the present. The extinction time of the species *s* born at time *t*_*c*_ is a random variable with a density distribution that is conditioned on extinction occurring before time *t*_*p*_,

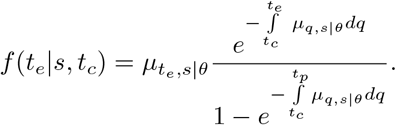

Note that this probability also depends on the extinction rate of the full process (i.e., at times later than *t*_*c*_), which is not always available, as it may depend, for example, on diversity at those later times. Hence, we propose to sample the extinction time from the truncated distribution

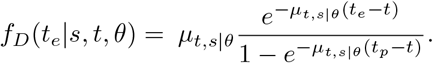

The full sampling probability of the missing part of a tree under this scheme is then given as

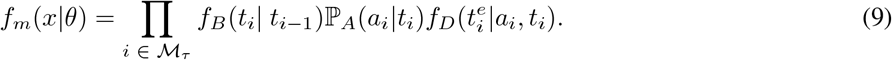

#### The data augmentation algorithm (DAA)

The main idea for our proposed data augmentation algorithm is to replace *P*_*α*_(*t*) by the probability that the newly created species (and not the entire clade that will descend from it) will become extinct before the present time

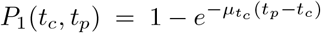

Thus, we consider the evolutionary process with diversification rates

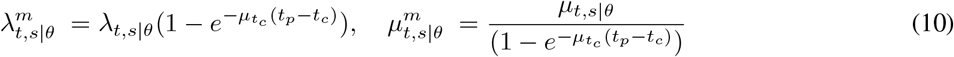

The algorithm is based on a Gillespie-type simulation algorithm which is a computationally simple and relatively simple digital computer algorithm [Gillespie, 1976, Kieu, 2018].

The algorithm proceeds as follows:

1. Input: Set *t*_0_ = 0, *i* = 1.
2. Draw a **missing speciation time** *t* from distribution

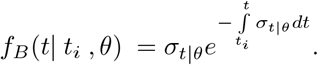

where

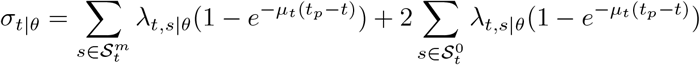
3. Draw an **allocation** for the species from distribution

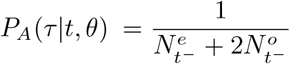
4. Draw the corresponding **extinction time** from distribution

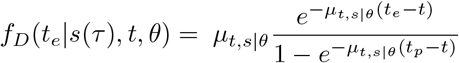
5. Set *t*_*i*_ = *t*, if *t*_*i*_ < *t*_*p*_ update the tree with the new species (speciation time, extinction time and allocation) and go to step 2; if *t*_*i*_ > *t*_*p*_ stop the algorithm and return the augmented tree.

An interpretation of this process is as follows:

- We observe a process with varying extinction rates, but as soon as a new missing species arises the extinction rate of that species is fixed throughout the rest of the process.
- When allocating a new species we assume that all possible allocations have a uniform probability distribution.
- We consider alternative probabilities *P*_*α*_(*t, T*), thus indexed by *α*, instead of the probability of not having any descendants *P*_0_(*t, T*). In our proposed importance sampler we consider the probability of extinction *P*_1_ of the just created lineage.

Figure 5 shows a diagram with the steps of the proposed data augmentation algorithm.

**Figure 5:**
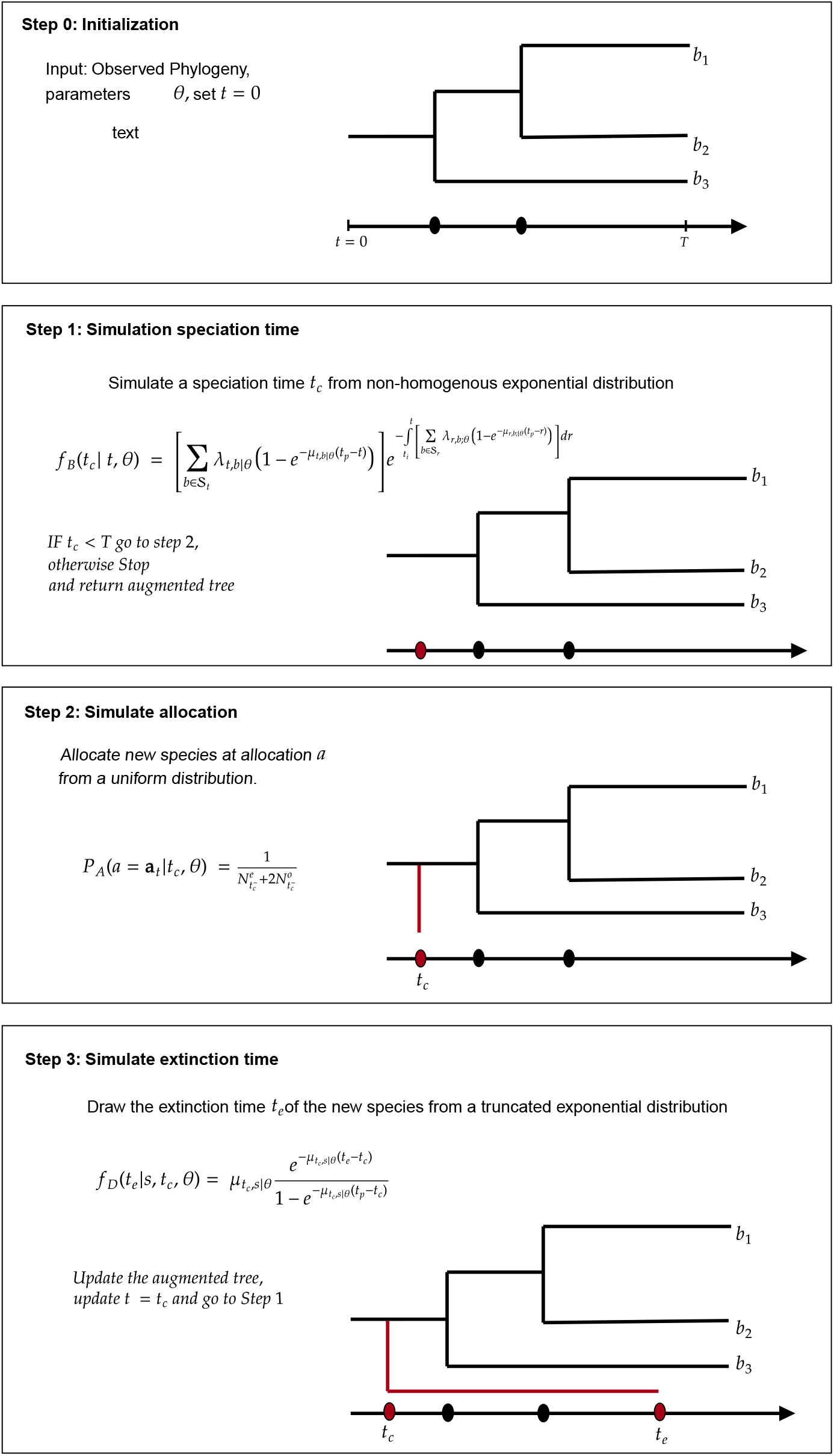
Tree augmentation algorithm based on the underlying non-homogeneous Poisson process.

Using equation (9) with the data augmentation scheme described above we have the following sampling probability of the full augmentation process:

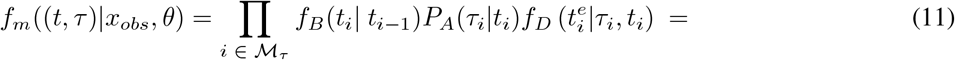

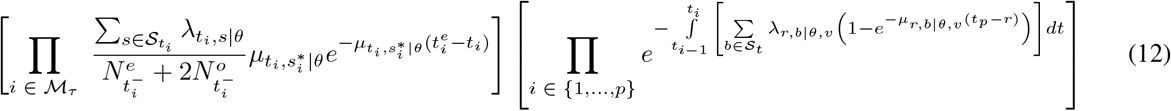

Taking the logarithm we have

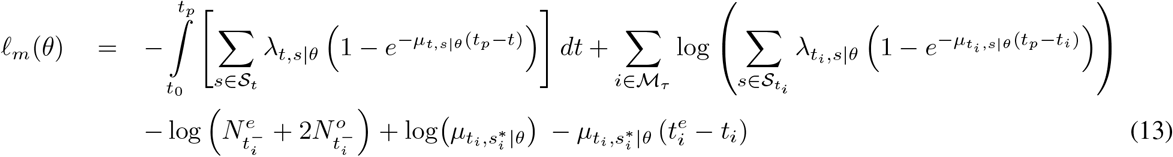

**Example**. Consider a model with a speciation rate that is the same for all lineages and with a constant extinction rate, i.e.,

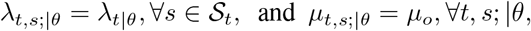

then, the sampling probability of the DAA is

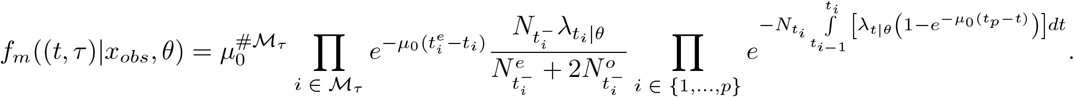

#### 3.3.1 Sample size

In Monte-Carlo methods, the variance of the estimates and the convergence time are determined by the sample size, the explored region of the parameter space and the type of data. From these three factors, we have control only over the sample size. MC methods require a sensible choice of the sample size, and it much depends on the type of problem. In iterative algorithms such as the MCEM algorithm, it is usually efficient to start with small sample size and increase it while parameters are approaching the MLE [Delyon et al., 1999], but there is no general rule for the choice of sampling sizes [Atanassov and Dimov, 2008].

To determine the required sample size in the emphasis method, we consider the estimator the distribution *f*_*m*_.

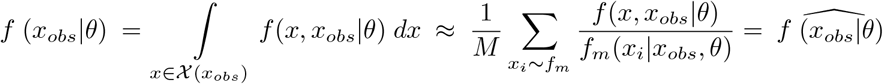

where {*x*_1_, …, *x*_*M*_} are full trees sampled from *f*_*m*_((*t, τ*)|*x*_*obs*_, *θ*). We will assume that if 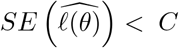 then, 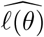 is good enough, for a small constant *C*. Note that, by taking a Taylor expansion of the logarithm of the estimated likelihood around the observed tree *x*_*obs*_, we can write

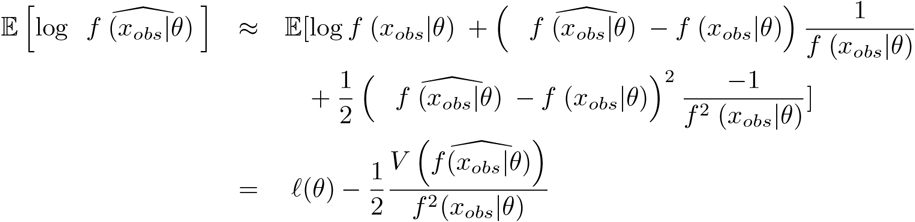

where the last term represents the first-order bias. So, typically our method will underestimate the loglikelihood. Furthermore, the estimation tends to be variable. The variability can be assessed by a first order Taylor expansion, i.e.,

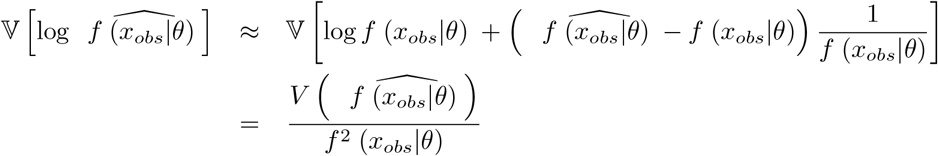

The variance of 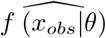 can be easily estimated by the sample variance of the importance weights divided by the sample size, i.e., 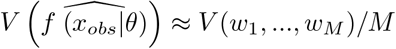. To assess the total possible deviation of the MC estimation we consider the bias and the standard error combined:

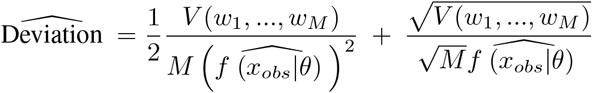

where the weights *w*_*i*_ are given by equation 6. The first term of the equation is an estimate of the bias and the second term an estimate of the standard error.

If it is feasible to perform a large number of trial simulations, then the standard error becomes of a lower order than the bias and, thus, we have that approximately,

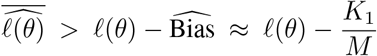

for a constant value 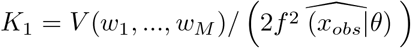, which can be further used to do a bias-correction of our likelihood. With this, we can calculate an approximation of the required sample size *M* to reach a desired level of accuracy. Figure 6 shows an illustration on the method we use to assess the required sample size. We sample trees with different sample sizes in order to obtain different estimates of the loglikelihood, and we can fit a curve of the form 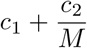. With the fitted model we can calculate the asymptotic value of the loglikelihood *c*_1_. We set the sample size *M* such that for a given tolerance level 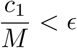.

**Figure 6:**
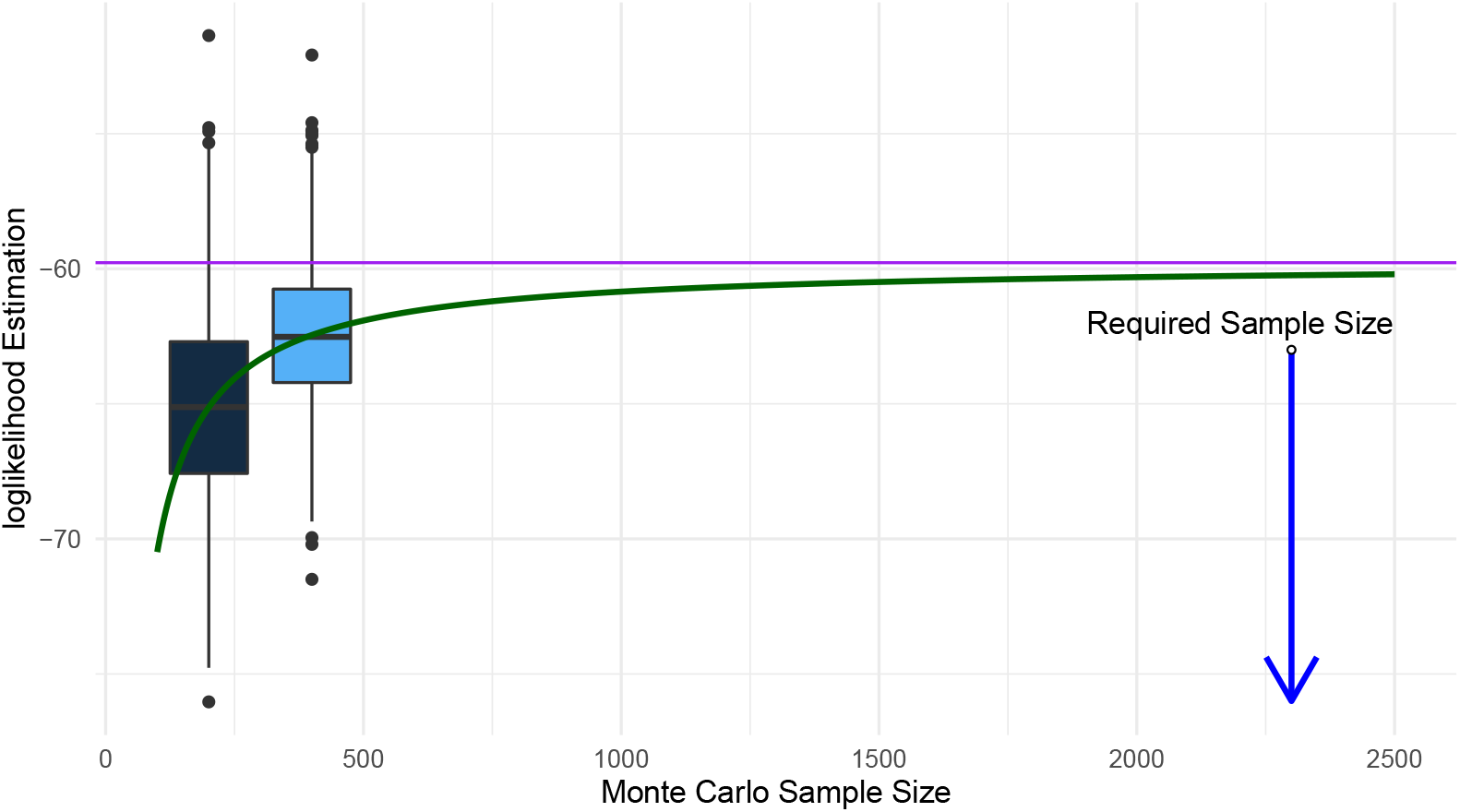
To calculate the required sample size, we simulate trees and estimate the loglikelihood via Monte-Carlo with at least two different sample sizes. We then calculate the curve that fits the relationship between the sample size and the estimated MC loglikelihood. This curve indicates the sample size required under a given tolerance level.

The MCEM algorithm can be replaced by the SAEM, MCMC or variations and combinations of them [Delyon et al., 1999, Celeux et al., 1995, Rydén et al., 2008, Wang, 2007, Kuhn and Lavielle, 2004]. All these algorithms rely on a sampling scheme and importance samplers. Our sampling scheme and sample size determination strategy can be used in any of these methods.

### 3.4 Model Selection

It is possible to apply standard model selection tools such as AIC or BIC [Wit et al., 2012] to the obtained loglikelihood. Furthermore, in the context of phylogenetic trees, specific statistics have been developed to test how well a model describes an observed tree. An informative summary statistic is the lineage-through-time (LTT) statistic [Janzen et al., 2015], defined as

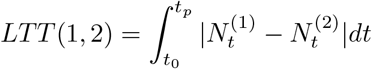

where 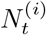 is the number of species of a tree *i* at time *t*. This statistic can also be used to assess how well a model describes an observed tree, simulating trees from the desired model and then calculating the LTT statistic between each simulated tree and the observed tree. It is also possible to calculate the mean number of species through time into a single “average” tree and calculate the LTT statistic of that tree compared to the observed tree.

The LTT statistic and model 1 are mathematical expressions that take into account the branching times of the tree, but ignore the topology. That is, the parent-child relationship among species is not relevant; only the branching times are considered. In this manuscript, we introduce an alternative to the LTT statistic, considering phylogenetic diversity instead of species richness. We define the *phylodiversity-through-time* (PTT) statistic as

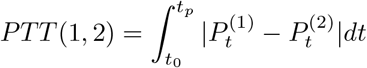

where 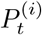 is the phylogenetic diversity for tree *i* at time *t*. In Figure 7 we present two example trees and the species richness for both trees as well as the phylogenetic diversity. The blue area represents the LTT statistic, while the green area represents the PTT statistic.

**Figure 7:**
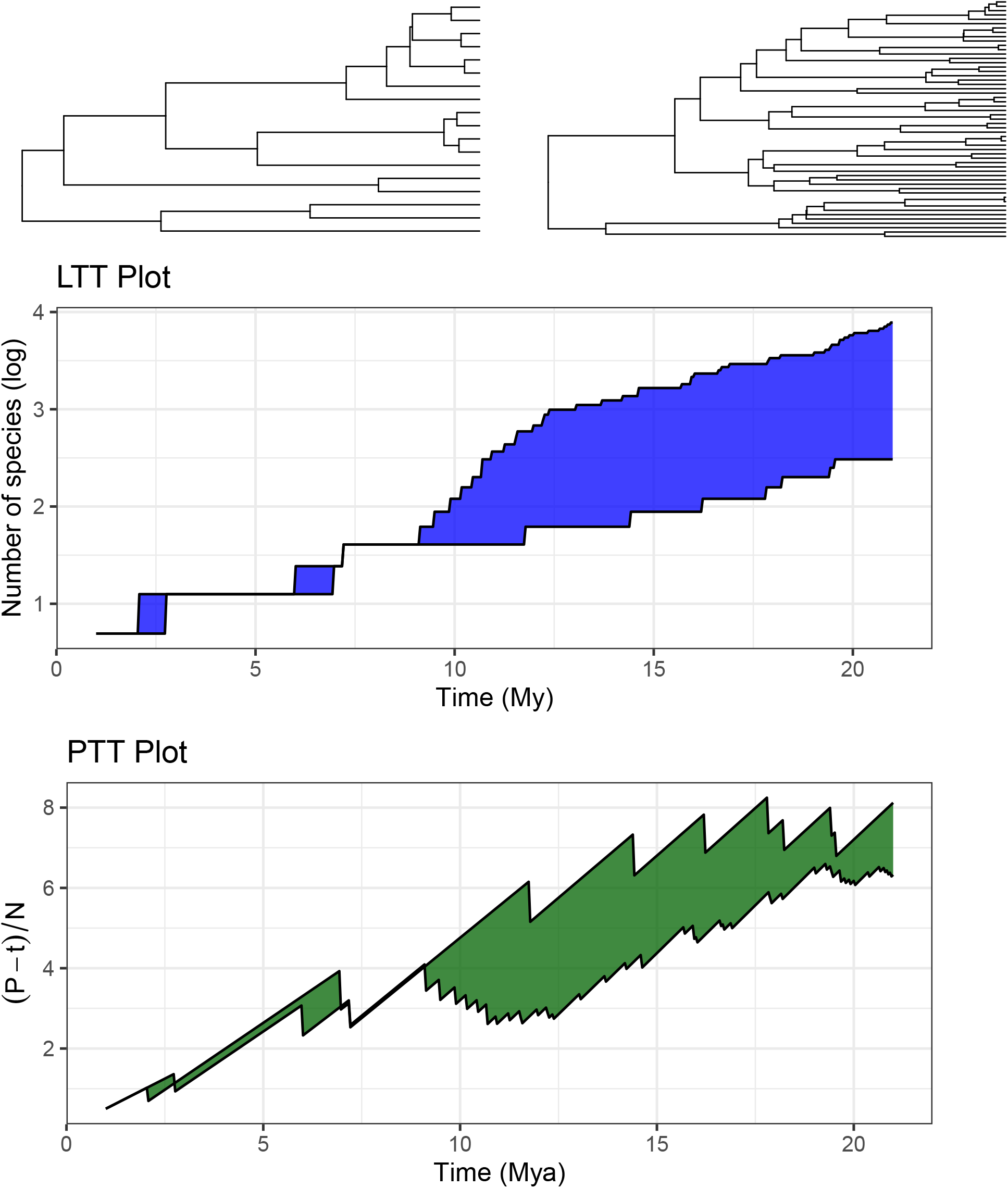
Comparison between two phylogenetic trees using the LTT (Number of lineages) and PTT (Phylogenetic diversity) through time. The area represents the distance between the trees.

Here, we will consider the LTT statistic, the PTT statistic and the AIC weights for model comparison and general goodness-of-fit considerations.

## 4 Application

To illustrate our method, we quantitatively compare model (2) with model (1) for 14 phylogenies obtained from [Condamine et al., 2019], with sizes ranging between 16 and 141 species and crown ages between 5 My and 65 My. Figure 8 represents the distribution of the number of species and crown age of the clades.

**Figure 8:**
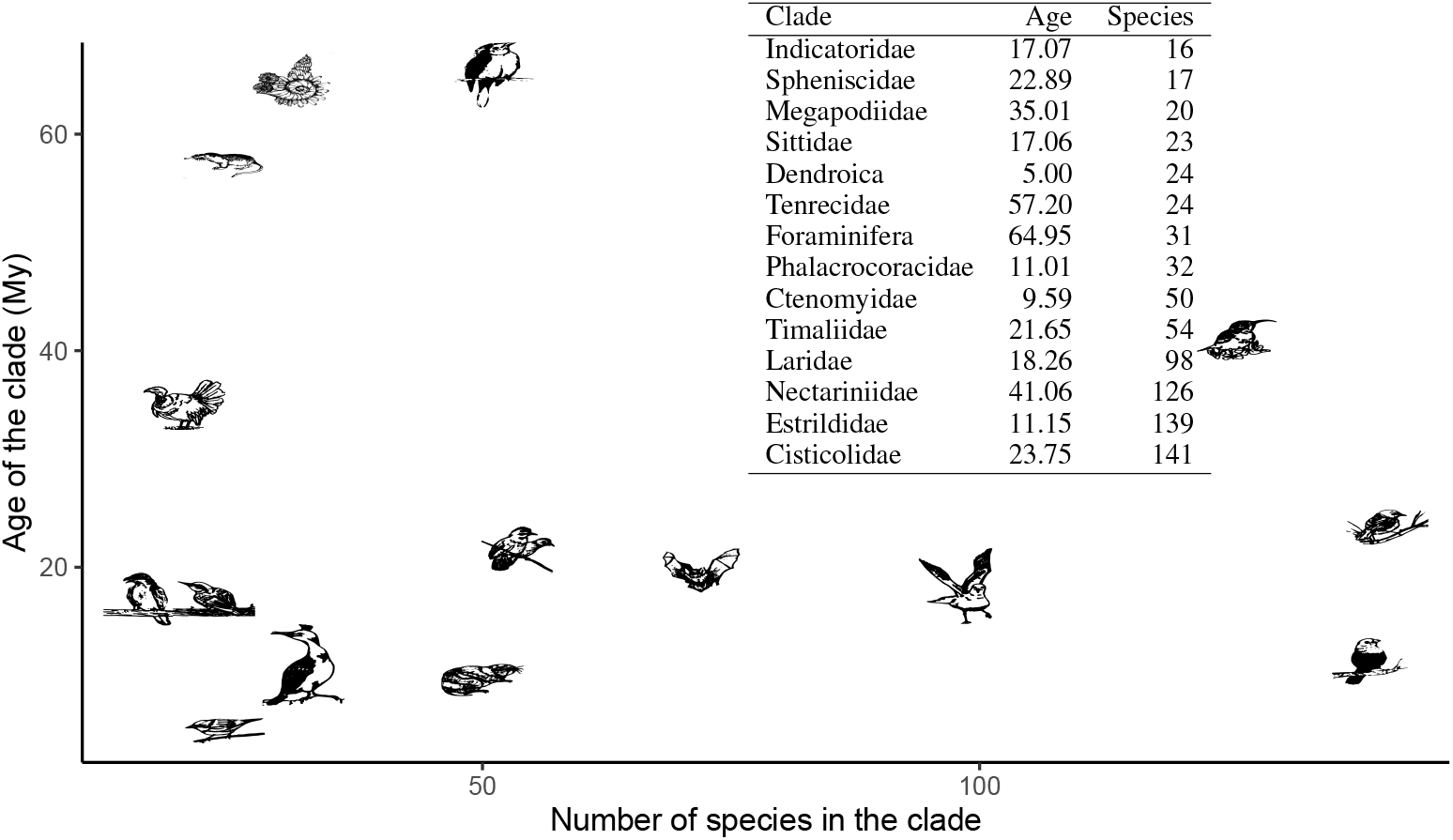
Distribution of crown ages and numbers of species of 14 phylogenetic trees.

In this application, the ultimate use of the emphasis framework is to quantify the impact of phylodiversity-dependent diversification by finding the maximum likelihood estimates for model (2) and compare them with model (1). But first, we perform some initial steps to evaluate the required sampling size for different phylogenetic trees. This will give insight about which phylogenies we can apply emphasis to and at what computational cost.

### 4.1 Monte-Carlo approximation with the proposed importance sampler

Before performing an analysis with the model (2), we want to test the efficiency of the Monte-Carlo method with the importance sampler introduced in Section 3.3. Monte-Carlo methods require a sensible choice of the sample size, and this largely depends on the type of problem. For sampling full trees, the relationship between accuracy and sample size is complex. For the uniform importance sampler presented in Richter et al. [2020], the required MC sample size becomes huge for most empirical trees. We first show that the non-homogenous sampler presented here can provide accurate approximations.

Note that,

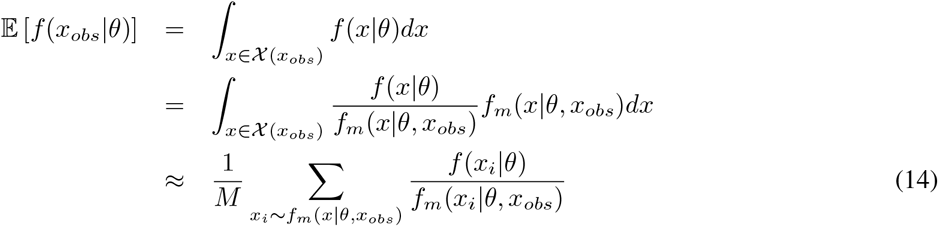

so we can use the Monte-Carlo sampling to approximate the likelihood for every parameter. To assess how well our importance sampler does as a function of MC sample size, we compare MC estimations for the 14 phylogenies for the LDD model, for which an existing solution exists. For each phylogeny, we calculate the MLE for the LDD model with the DDD R package. With these parameters, we perform Monte-Carlo sampling and approximate the expectation (14) with 4 different MC sampling sizes. In Table 1 we show the MC estimations and, in the last column, the analytical solution.

**Table 1:** Monte-Carlo approximation of the loglikelihood for each tree at its corresponding MLE for the diversification process generated under the LDD model, for different sample sizes. The last column contains the analytical value obtained with the R package DDD.

| | $10^2$ | $10^3$ | $10^4$ | $10^5$ | $10^6$ | Analytical |
| --- | --- | --- | --- | --- | --- | --- |
| Indicatoridae | -42.04(1.1e-01) | -42.39(1.1e-01) | -41.97(4.9e-02) | -42.01(4.1e-02) | -41.95(4.8e-02) | -41.89 |
| Spheniscidae | -50.22(3.4e-01) | -50.64(1.8e-01) | -50.49(1.1e-01) | -50.23(8.1e-02) | -50.35(4.3e-02) | -50.23 |
| Megapodiidae | -68.98(6.1e-02) | -68.77(4.6e-02) | -68.3(1.9e-02) | -68.1(2.1e-02) | -68.09(1.0e-02) | -68 |
| Sittidae | -64.72(3.8e-01) | -64.19(2.3e-01) | -64.42(1.0e-01) | -64.38(7.6e-02) | -64.32(3.0e-02) | -64.31 |
| dendroica | -38.97(3.5e-01) | -39.2(2.3e-01) | -39.04(1.1e-01) | -38.97(6.0e-02) | -39.03(4.3e-02) | -38.91 |
| Tenrecidae | -89.68(1.7e-01) | -89.12(6.0e-02) | -88.84(3.6e-02) | -89.05(2.1e-02) | -88.42(4.4e-02) | -88.4 |
| foraminifera | -118.48(5.9e-02) | -117.7(4.3e-02) | -116.48(2.4e-02) | -116.34(1.9e-02) | -115.75(1.1e-02) | -115.73 |
| Phalacrocoracidae | -79.92(4.6e-02) | -81.06(9.5e-02) | -80.42(5.7e-02) | -80.46(4.0e-02) | -80.4(2.0e-02) | -80.25 |
| Ctenomyidae | -122.66(1.4e-01) | -120.8(1.1e-01) | -120.61(6.1e-02) | -120.7(4.0e-02) | -120.63(5.6e-02) | -120.65 |
| Timaliidae | -154.23(4.7e-03) | -154.93(5.1e-03) | -153.75(3.0e-03) | -153.91(2.2e-03) | -153.56(9.8e-04) | -153.48 |
| Laridae | -232.12(5.9e-03) | -232.15(3.1e-03) | -231.52(2.0e-03) | -231.23(1.1e-03) | -231.14(8.7e-04) | -231.07 |
| Nectariniidae | -416.2(6.3e-04) | -410.81(2.7e-05) | -404.61(2.0e-06) | -402.19(7.4e-07) | -400.74(3.3e-07) | -399.04 |
| Estrildidae | -321.31(5.4e-03) | -316.31(1.7e-04) | -311.26(2.7e-05) | -312.22(2.8e-05) | -310.93(1.4e-05) | -309.35 |
| Cisticolidae | -410.16(2.5e-04) | -403.73(2.3e-05) | -402.29(1.0e-05) | -402.06(7.3e-06) | -400.6(4.0e-06) | -397.94 |

If the difference between the analytical loglikelihood and the MC approximated loglikelihood is less than 1, we will conclude that the estimation is good enough, following the AIC principle when comparing a model with three parameters agains a model with four parameters. Under this assumption, we found that a sample size of 1000 is good enough for phylogenies up to approximately 70 species. With a sample size of 10^6^ we have very accurate estimations for all clades with the exceptions of Nectariniidae, Estrildidae and Cisticolidae. These are the larger trees with more than 100 species. Table 1 contains detailed estimation for 4 different sample sizes for each phylogeny.

### 4.2 Estimation and model selection

In Table 2 we report the parameter estimations for the two models of interest for the 14 case-study phylogenies. Looking at the LTT statistics, we observe that there is only a slight improvement of the LPDD model over the LDD model in most of the cases. This is confirmed by the loglikelihood values, where the improvement is not large enough to justify preferring the LPDD model, which is confirmed with the AIC weights which always prefer the LDD model.

**Table 2:** Parameter estimations for LDD and LPDD model for 14 phylogenies. The fifth column shows the AIC weights for the comparison of these two models. The sixth column is the normalised LTT statistic. The last four columns represent the parameter estimates. Between parentheses we report the standard deviation of the Monte-Carlo approximation.

| Clade | Age | Tips | Model | AICw | LTT | PTT | loglikelihood | $\mu_0$ | $\lambda_0$ | $\beta_N$ | $\beta_P$ |
| --- | --- | --- | --- | --- | --- | --- | --- | --- | --- | --- | --- |
| Indicatoridae | 17.07 | 16 | LDD | 0.73 | 0.49 | 0.49 | -42.01 | 0.22 | 1.62 | -0.085 | 0 |
|  |  |  | LPDD | 0.27 | 0.51 | 0.51 | -41.99 (6e-02) | 0.21 (4e-03) | 1.61 (5e-04) | -0.085 (5e-05) | -0.001 (3e-04) |
| Spheniscidae | 22.89 | 17 | LDD | 0.83 | 0.41 | 0.52 | -50.23 | 0.2 | 1.61 | -0.081 | 0 |
|  |  |  | LPDD | 0.17 | 0.59 | 0.48 | -50.79 (3e-02) | 0.16 (2e-03) | 1.49 (1e-02) | -0.079 (6e-04) | 0.004 (7e-04) |
| Megapodiidae | 35.01 | 20 | LDD | 0.78 | 0.45 | 0.48 | -68.1 | 0.1 | 0.83 | -0.036 | 0 |
|  |  |  | LPDD | 0.22 | 0.55 | 0.52 | -68.37 (3e-02) | 0.09 (1e-03) | 0.83 (6e-04) | -0.036 (4e-05) | 0 (1e-04) |
| Sittidae | 17.06 | 23 | LDD | 0.79 | 0.5 | 0.46 | -64.38 | 0.15 | 0.58 | -0.018 | 0 |
|  |  |  | LPDD | 0.21 | 0.5 | 0.54 | -64.72 (1e-01) | 0.12 (9e-04) | 0.4 (1e-03) | -0.021 (3e-04) | 0.039 (1e-03) |
| Dendroica | 5.00 | 24 | LDD | 0.77 | 0.46 | 0.53 | -38.97 | 0.16 | 3.05 | -0.117 | 0 |
|  |  |  | LPDD | 0.23 | 0.54 | 0.47 | -39.16 (8e-02) | 0.14 (1e-03) | 2.99 (2e-02) | -0.118 (1e-03) | 0.007 (3e-03) |
| Tenrecidae | 57.20 | 24 | LDD | 0.74 | 0.46 | 0.47 | -89.05 | 0.11 | 0.59 | -0.02 | 0 |
|  |  |  | LPDD | 0.26 | 0.54 | 0.53 | -89.1 (3e-02) | 0.09 (6e-04) | 0.59 (4e-04) | -0.02 (8e-05) | 0.001 (2e-04) |
| Foraminifera | 64.95 | 31 | LDD | 0.84 | 0.39 | 0.42 | -116.34 | 0.1 | 1.18 | -0.034 | 0 |
|  |  |  | LPDD | 0.16 | 0.61 | 0.58 | -117.01 (4e-03) | 0.08 (4e-04) | 1.17 (1e-04) | -0.034 (2e-06) | 0 (4e-06) |
| Phalacrocoracidae | 11.01 | 32 | LDD | 0.71 | 0.41 | 0.48 | -80.46 | 0.24 | 1.67 | -0.044 | 0 |
|  |  |  | LPDD | 0.29 | 0.59 | 0.52 | -80.34 (4e-02) | 0.24 (3e-03) | 1.59 (2e-02) | -0.038 (3e-04) | -0.027 (3e-03) |
| Ctenomyidae | 9.59 | 50 | LDD | 0.74 | 0.46 | 0.42 | -120.7 | 0.16 | 1.15 | -0.02 | 0 |
|  |  |  | LPDD | 0.26 | 0.54 | 0.58 | -120.75 (6e-02) | 0.14 (1e-03) | 1.08 (6e-03) | -0.019 (2e-04) | 0.007 (2e-03) |
| Timaliidae | 21.65 | 54 | LDD | 1 | 0.48 | 0.53 | -153.91 | 0.14 | 0.5 | -0.006 | 0 |
|  |  |  | LPDD | 0 | 0.52 | 0.47 | -158.63 (7e-03) | 0.07 (1e-03) | 0.22 (8e-03) | -0.006 (2e-04) | 0.034 (2e-03) |
| Laridae | 18.26 | 98 | LDD | 0.68 | 0.19 | 0.5 | -231.23 | 0.13 | 0.32 | 0 | 0 |
|  |  |  | LPDD | 0.32 | 0.81 | 0.5 | -231.01 (6e-02) | 0.02 (1e-03) | 0.46 (2e-03) | 0.001 (2e-05) | -0.061 (7e-04) |
| Nectariniidae | 41.06 | 126 | LDD | 0.66 | 0.54 | 0.83 | -402.19 | 0.14 | 0.32 | -0.001 | 0 |
|  |  |  | LPDD | 0.34 | 0.46 | 0.17 | -401.83 (4e-02) | 0.02 (4e-04) | 0.25 (1e-03) | 0 (2e-05) | -0.019 (3e-04) |
| Estrildidae | 11.15 | 139 | LDD | 0.88 | 0.5 | 0.77 | -312.22 | 0.28 | 1.05 | -0.005 | 0 |
|  |  |  | LPDD | 0.12 | 0.5 | 0.23 | -313.21 (6e-03) | 0.12 (2e-03) | 0.42 (7e-03) | -0.006 (3e-04) | 0.176 (1e-02) |
| Cisticolidae | 23.75 | 141 | LDD | 0.51 | 0.48 | 0.76 | -402.06 | 0.16 | 0.48 | -0.002 | 0 |
|  |  |  | LPDD | 0.49 | 0.52 | 0.24 | -401.11 (2e-02) | 0.05 (1e-03) | 0.37 (4e-03) | 0 (5e-05) | -0.04 (1e-03) |

Note that the AIC weights are based on a Monte-Carlo approximation of the likelihood which will slightly underestimate models with more parameters. As a result, the AIC is more conservative than a test where the AIC values are calculated using the true value of the loglikelihood. Note that in some cases the loglikelihood for the LPDD model is still smaller than the loglikelihood of the LDD model, which cannot be correct because the LDD model is nested within the LPDD model, and hence the LPDD likelihood should always be smaller than the LPDD likelihood. We argue that this is because the Monte-Carlo approximation is not good enough yet. These computational issues suggest that hypothesis testing with AIC might not be an appropriate tool for model selection. Significance tests, instead, do not depend on the approximation of the likelihood but on the approximation of the Hessian of the likelihood (see equation **??**); because the likelihood is asymptotically quadratic near its maximum (hence the second derivative is constant), the approximation of the Hessian should not present the computational issues that the approximation of the likelihood presents, and hence significance tests seem more reliable. Based on the significance test results, we conclude that the phylodiversity-dependent diversification model provides an alternative/better explanation to/than the diversity-dependent diversification model, at least in some of our clades.

In Figure 9 we see an example of the expected lineages-through-time plot for each model in comparison with the observed lineage through time plot corresponding to the Timaliidae phylogeny, and the speciation rates through time plot. We can see that both models agree that speciation happened roughly at a rate of 0.2 species per million rate during the last 10 million years; however, they diverge on the estimates for the period between 20 and 10 million years ago. Including phylogenetic diversity involves a fluctuating speciation rate around 0.2 spe/Mye reaching its maximum around 15 years ago while the LDD model assumes a monotonously decreasing speciation rate. In general, the difference between the two models is not large and this pattern is present across all the 14 phylogenies.

**Figure 9:**
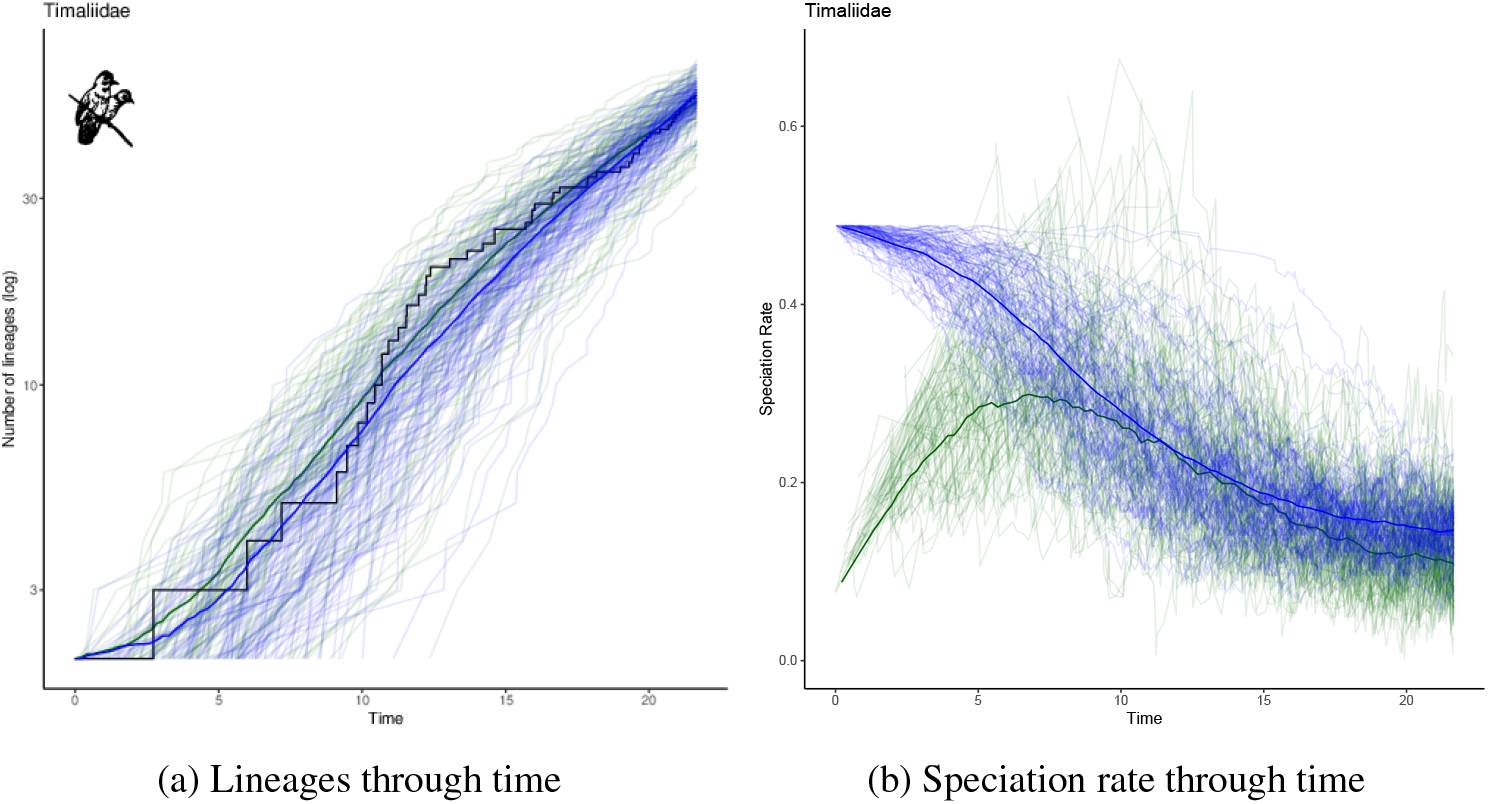
Evolution of extant species richness (LTT-plot) and evolution of global speciation rates for 13 clades under LDD (blue) and LPD (green) models.

Finally, in Table 3 we report the loglikelihood estimates for the LPDD model. We see that for most of the cases the sample size was large enough, but for larger trees the convergence performs much slower for the LPDD case than for the LDD case.

**Table 3:** Loglikelihood approximations of the LPD model at its MLE value for the 14 phylogenies.

| | $10^2$ | $10^3$ | $10^4$ | $10^5$ | $10^6$ |
| --- | --- | --- | --- | --- | --- |
| Indicatoridae | -42.49(1.3e-01) | -42.15(1.0e-01) | -42.26(6.5e-02) | -41.99(6.1e-02) | -42.02(2.7e-02) |
| Spheniscidae | -51.66(2.4e-01) | -50.68(1.1e-01) | -50.9(6.4e-02) | -50.79(3.0e-02) | -50.75(1.6e-02) |
| Megapodiidae | -68.97(2.1e-01) | -68.19(8.1e-02) | -68.42(6.2e-02) | -68.37(3.5e-02) | -68.26(4.0e-02) |
| Sittidae | -65.49(2.7e-01) | -64.49(2.0e-01) | -64.83(1.2e-01) | -64.72(1.2e-01) | -64.73(2.4e-02) |
| dendroica | -39.25(2.9e-01) | -39.33(1.9e-01) | -39.22(9.4e-02) | -39.16(8.0e-02) | -39.14(3.2e-02) |
| Tenrecidae | -89.06(8.6e-02) | -88.65(6.0e-02) | -89.61(3.9e-02) | -89.1(3.4e-02) | -89.12(1.5e-02) |
| foraminifera | -119.3(1.2e-02) | -117.71(5.1e-03) | -117.94(3.4e-03) | -117.01(4.5e-03) | -117.3(2.3e-03) |
| Phalacrocoracidae | -79.61(1.2e-01) | -81.02(8.5e-02) | -80.54(6.5e-02) | -80.34(4.1e-02) | -80.56(2.1e-02) |
| Ctenomyidae | -119.95(1.8e-01) | -121.08(2.0e-01) | -120.87(8.2e-02) | -120.75(5.6e-02) | -120.77(7.6e-02) |
| Timaliidae | -159.57(4.8e-02) | -158.95(2.0e-02) | -158.37(1.4e-02) | -158.63(6.8e-03) | -158.16(6.9e-03) |
| Laridae | -230.98(6.3e-01) | -230.86(2.2e-01) | -231.03(1.0e-01) | -231.01(5.6e-02) | -231(3.8e-02) |
| Nectariniidae | -402.09(1.9e-01) | -402.01(9.6e-02) | -401.81(6.9e-02) | -401.83(3.9e-02) | -401.87(3.0e-02) |
| Estrildidae | -315.33(3.0e-02) | -313.25(1.3e-02) | -313.13(1.1e-02) | -313.21(6.0e-03) | -313.28(3.7e-03) |
| Cisticolidae | -403.78(5.9e-02) | -401.9(3.6e-02) | -401.52(2.4e-02) | -401.11(1.7e-02) | -401(8.2e-03) |

## 5 Discussion

Diversity-dependent diversification models have been developed during the last decade in order to understand and quantify the existence and impact of ecological limits to macroevolutionary dynamics. At the moment, only models with a dependence of diversification rates on species richness have been implemented, but these models ignore other facets of diversity, such as phylodiversity.

Here, we have completed the statistical methodology introduced in Richter et al. [2020], with the design of a data augmentation scheme that provides an efficient importance sampler. This is a substantial improvement in comparison to the uniform importance sampler considered in Richter et al. [2020], as it enables applying the method to a large number of empirical phylogenies.

In the application to 14 example phylogenies, we studied the LPDD model, i.e., a model with a linear effect of phylodiversity on speciation. We found that including phylodiversity does not provide a substantial improvement in comparison with richness-dependent diversification models. However, phylodiversity does provide an alternative and slightly more complete explanation to speciation dynamics; the LTT statistic and the PTT statistic provide insights and, most of the times, reflect that trees generated by the LPDD model are closer to real phylogenies than trees generated under the LDD model. While the model with fewer parameters is preferred using AIC, the phylodiversity component is statistically significant, suggesting that it should not be ignored.

This may not be the final word because there are some technical improvements to be made. In particular, we did not condition the likelihood on non-extinction of the clade; even though this is generally recommended [Etienne et al., 2016, Stadler, 2013].

Our method is not limited to phylogenetic diversity-dependent diversification models, but allows inference of a general class of species diversification models, considering time, traits, climate, functional diversity, just to name a few. With the data augmentation described her we have provided a general tool that can be potentially used to quantify and test a large number of hypotheses in macroevolutionary diversification.

## 6 Acknowledgements

We thank Alexei Drummond for helpful comments on this manuscript and Bart Haegeman for insightful conversations about diversity-dependence models.

This publication is part of the project *Killing two birds with one stone: simultaneous estimation and selection of species diversification models* with project number 657.014.005 of the research programme *Mathematics for planet Earth (MPE)* which is partly financed by the Dutch Research Council (NWO).

This article is based upon work from COST Action *European Cooperation for Statistics of Network Data Science* (CA15109), supported by COST (European Cooperation in Science and Technology).

We also acknowledge funding from the Swiss National Science Foundation, project 200021_188534 entitled Sparse inference of complex networks.

## 7 Author contributions

F.R., E.W and R.E. conceived of the presented idea and developed the theoretical formalism. F.R, T.J. and H.H. carried out the implementation. F.R. performed the computations. F.R, E.W and R.E wrote the manuscript. F.R, E.W, R.E and T.J discussed the results and contributed to the final manuscript. E.W. and R.E supervised the project.

